# Looking into Pandora’s Box: The Content of *Sci-Hub* and its Usage

**DOI:** 10.1101/124495

**Authors:** Bastian Greshake

**Affiliations:** Institute of Cell Biology and Neuroscience, Goethe University Frankfurt, Frankfurt, Germany

## Abstract

Despite the growth of Open Access, illegally circumventing paywalls to access scholarly publications is becoming a more mainstream phenomenon. The web service *Sci-Hub* is amongst the biggest facilitators of this, offering free access to around 62 million publications. So far it is not well studied how and why its users are accessing publications through *Sci-Hub*. By utilizing the recently released corpus of *Sci-Hub* and comparing it to the data of ˜28 million downloads done through the service, this study tries to address some of these questions. The comparative analysis shows that both the usage and complete corpus is largely made up of recently published articles, with users disproportionately favoring newer articles and 35% of downloaded articles being published after 2013. These results hint that embargo periods before publications become Open Access are frequently circumnavigated using Guerilla Open Access approaches like *Sci-Hub*. On a journal level, the downloads show a bias towards some scholarly disciplines, especially Chemistry, suggesting increased barriers to access for these. Comparing the use and corpus on a publisher level, it becomes clear that only 11% of publishers are highly requested in comparison to the baseline frequency, while 45% of all publishers are significantly less accessed than expected. Despite this, the oligopoly of publishers is even more remarkable on the level of content consumption, with 80% of all downloads being published through only 9 publishers. All of this suggests that *Sci-Hub* is used by different populations and for a number of different reasons, and that there is still a lack of access to the published scientific record. A further analysis of these openly available data resources will undoubtedly be valuable for the investigation of academic publishing.

## Introduction

Through the course of the 20th century, the academic publishing market has radically transformed. What used to be a small, decentralized marketplace, occupied by university presses and educational publishers, is now a global, highly profitable enterprise, dominated by commercial publishers [1]. This development is seen as the outcome of a multifactorial process, with the inability of libraries to resist price increases, the passivity of researchers who are not directly bearing the costs and the merging of publishing companies, leading to an oligopoly [2].

In response to these developments and rising subscription costs, the Open Access movement started out to reclaim the process of academic publishing [3]. Besides the academic and economic impact, the potential societal impact of Open Access publishing is getting more attention, [4, 5] and large funding bodies seem to agree with this opinion, as more and more are adopting Open Access policies [6, 7, 8]. These efforts seem to have an impact, as a 2014 study of scholarly publishing in the English language found that, while the adoption of Open Access varies between scholarly disciplines, an average of around 24 % of scholarly documents are freely accessible on the web [9]. Another response to these shifts in the academic publishing world is what has been termed *Guerilla Open Access* [1], *Bibliogifts* [10] or *Black Open Access* [11]. Or in short, the usage of semi-legal or outright illegal ways of accessing scholarly publications, like peer2peer file sharing, for example the use of *#icanhazpdf* on Twitter [10], or centralized web services like *Sci-Hub/LibGen* [12].

Especially *Sci-Hub*, which started in 2011, has moved into the spotlight in the recent years. According to founder Alexandra Elbakyan, the website uses donated library credentials of contributors to circumvent publishers’ paywalls and thus downloads large parts of their collections [13]. This clear violation of copyright not only lead to a lawsuit by Elsevier against Elbakyan [14], but also to her being called “*the Robin Hood of Science*” [15], with both sparking further interest in *Sci-Hub*.

Despite this, there has been little research into how *Sci-Hub* is used and what kind of materials are being accessed through it. A 2014 study has looked at content provided through *LibGen* [10]. In 2016 *Sci-Hub* released data on ∼ 28 million downloads done through the service [16]. This data was subsequently analyzed to see in which countries the website is being used, which publishers are most frequent [13] and how downloading publications through *Sci-Hub* relates to socio-economic factors, such as being based in a research institution [17] and how it impacts interlibrary loans [12].

In March 2017 *Sci-Hub* released the list of ∼ 62 million Digital Object Identifiers (DOIs) of the content they have stored. This study is the first to utilize both the data on which publications are downloaded through *Sci-Hub*, as well as the complete corpus available through them. This allows a data-driven approach to evaluate what is stored in the *Sci-Hub* universe, how the actual use of the service differs from that, and what different use cases people might have for *Sci-Hub*.

## Methods

### Data sources

The data on the around 62 million DOIs indexed by *Sci‐Hub* was taken from the dataset released on 2017-03-19 [18]. In addition, the data on the 28 million downloads done through *Sci-Hub* between September 2015 and February 2016 [16] was matched to the complete corpus of DOIs. This made it possible to quantify how often each object listed in *Sci-Hub* was actually requested from its user base.

### Resolving DOIs

The corresponding information for the publisher, the year of publication, as well as the journal in which it was published was gotten from doi.org, using the *RubyGem Terrier* (v1.0.2, https://github.com/Authorea/terrier). Acquiring the metadata for each of the 62 million DOIs in Sci-Hub was done between 2017-03-20 and 2017-03-31. In order to save time, the DOIs of the 28 million downloads were then matched to the superset of the already resolved DOI of the complete *Sci-Hub* catalog. In both cases, DOIs that could not be resolved were excluded from further analysis, but they are included in the dataset released with this article.

### Tests for over‐ & under-representation

For each publisher, the number of papers downloaded was compared to the expected number of downloads, given the publishers’ presence in the whole *Sci-Hub* database. For this the relative contribution to the database was calculated for each publisher, excluding all missing data. The number of actual downloads was then compared to the expected number of downloads using a binomial test. All p-values were corrected for multiple testing with False Discovery Rate [19] and post-correction p<0.05 were accepted.

## Results

### Resolving the *Sci-Hub* DOIs

For the 61,940,926 DOIs listed in the Sci-Hub data dump, a total of 46,931,934 DOIs could be resolved (75.77%). Manual inspection of the unresolvable 25% shows that nearly all of these could not be resolved as they are not available via doi.org, and are not a technical error in the procedure to resolve them (i.e. lack of internet connection). For the data on the downloads done through Sci-Hub, 21,515,195 downloads could be resolved out of 27,819,965 total downloads (77.34%).

### The age of publications in *Sci-Hub*

To estimate the age distribution of the publications listed in *Sci-Hub*, and which fraction of these publications is actually requested by the people using *Sci-Hub*, the respective datasets were tabulated according to the year of publication, see Figure 1. While over 95% of the publications listed in *Sci-Hub* were published after 1950, thereis nevertheless a long tail, reaching back to the 1619 edition of *Descriptio cometæ* [20].

**Figure 1.**
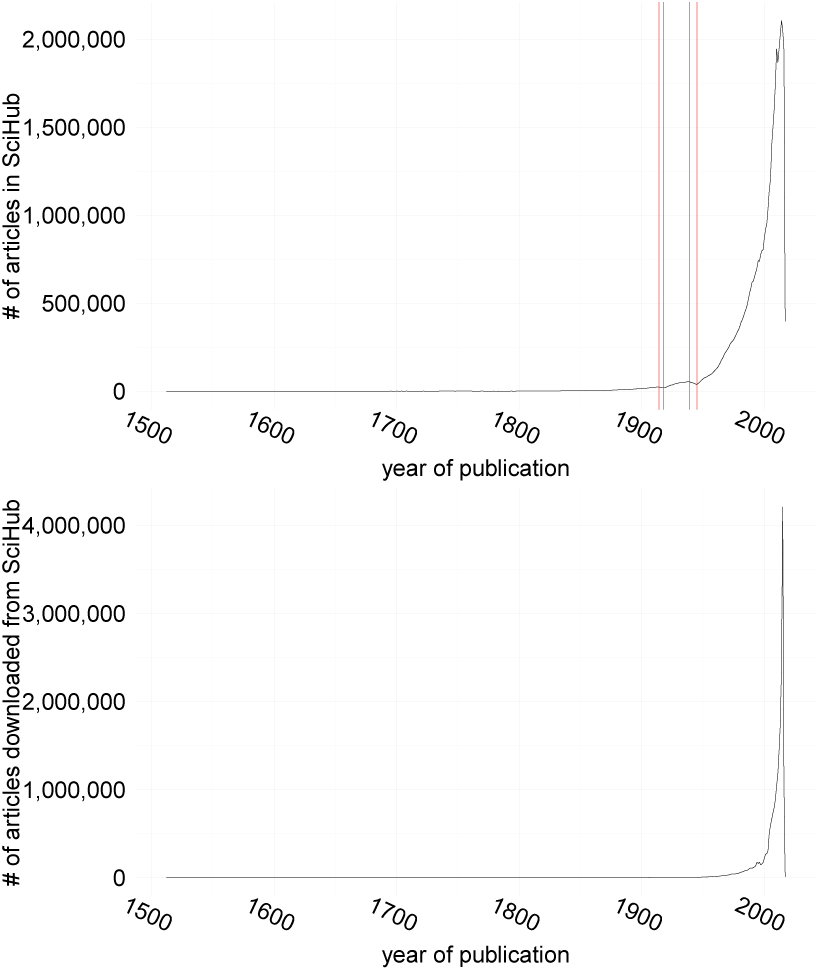
Top: Number of Publications in *Sci-Hub* by year of publication. Red bars denote the years 1914, 1918, 1939 and 1945. Bottom: Number of publications downloaded by year of publication.

As a general trend the number of publications listed in *Sci-Hub* increases from year to year. Two notable exceptions are the time periods of the two World Wars, at which ends the number of publications dropped to pre-1906 and pre-1926 levels, respectively (red bars in Figure 1).

When it comes to the publications downloaded by *Sci-Hub* users, the skew towards recent publications is even more extreme. Over 95% of all downloads fall into publications done after 1982, with ∼35% of the downloaded publications being less than 2 years old at the time they are being accessed (i.e. published after 2013). Despite this, there is also a long tail of publications being accessed, with articles published even in the 1600s being amongst the downloads, and 0.04% of all downloads being made for publications released prior to 1900.

### Which journals are being read?

The complete released database contains ∼177,000 journals, with ∼60% of these having at least a single paper downloaded. The number of articles per journal likely follows an exponential function, for both the total number of publications listed on *Sci-Hub* as well as the number of downloaded articles (see Supplementary Figure S1), with <10% of the journals being responsible for >50% of the total content in *Sci-Hub*. The skew for the downloaded content is even more extreme, with <1% of all journals getting over 50% of all downloads.

Contrasting the 20 most frequent journals in the complete database with the 20 most downloaded ones (Figure 2), one observes a clear shift not only in the distribution but also in the ranking, with the most abundant journal of the whole corpus not appearing in the 20 most downloaded journals. In addition, chemical journals appear to be overrepresented in the downloads (12 journals), compared to the complete corpus (7 journals), with no other discipline showing an increase amongst the 20 most frequent journals.

**Figure 2.**
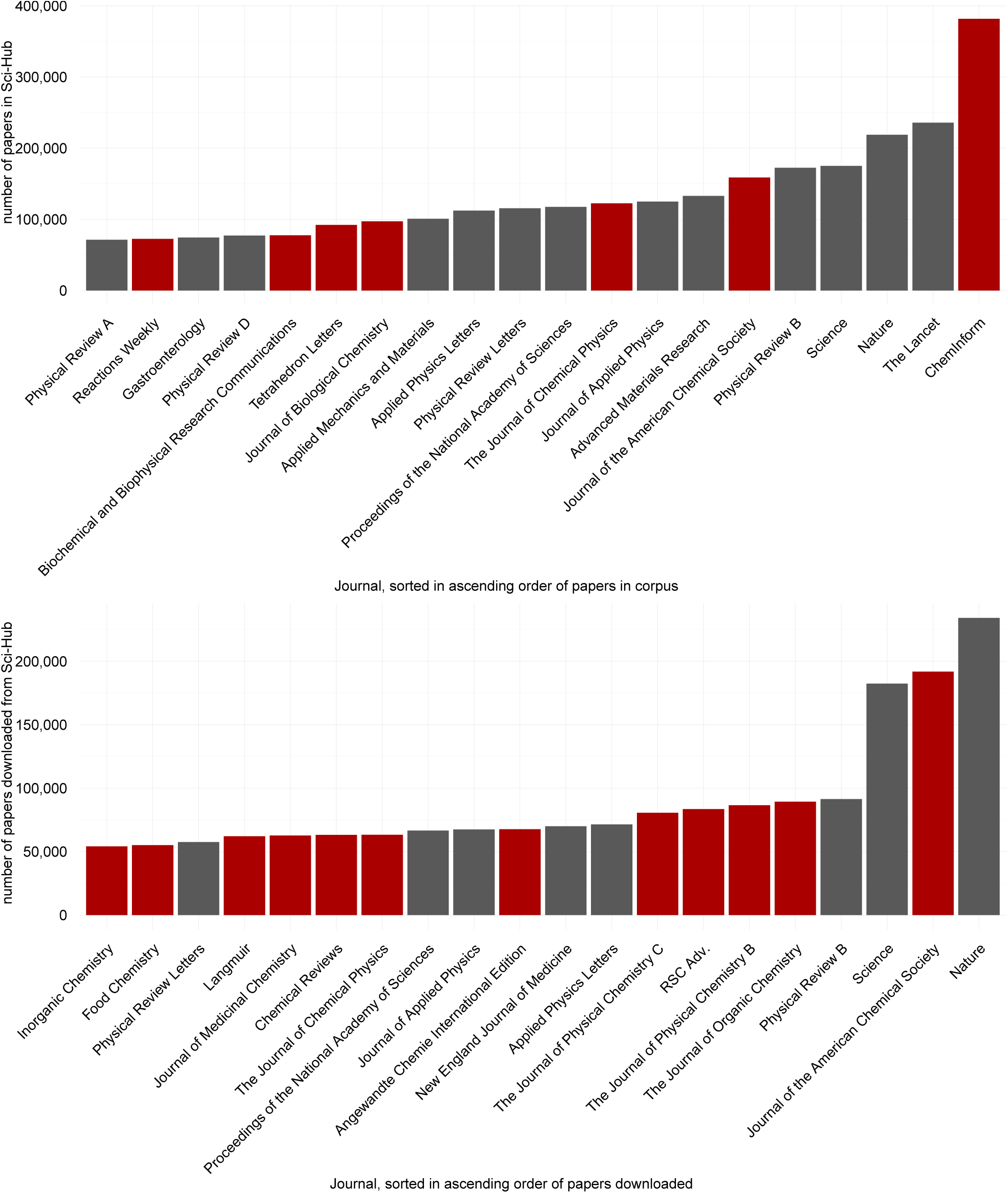
Top: The 20 most frequent journals in all of *Sci-Hub*. Bottom: The 20 journals with the most downloads. In both panels Chemistry journals are highlighted in red.

### Are publishers created equal?

Looking at the data on a publisher level, there are ∼1,700 different publishers, with ∼1,000 having at least a single paper downloaded. Both corpus and downloaded publications are heavily skewed towards a set of few publishers, with the 9 most abundant publishers having published ∼70% of the complete corpus and ∼80% of all downloads respectively (see Supplementary Figure S2). Given the background frequency in the complete corpus, the download numbers were compared to the expected numbers using a binomial test. After false discovery rate correction for multiple testing, 982 publishers differed significantly from the expected download numbers, with 201 publishers having more downloads than expected and 781 being underrepresented. Interestingly, while some big publishers like Elsevier and Springer Nature come in amongst the overly downloaded publishers, many of the large publishers, like Wiley-Blackwell and the Institute of Electrical and Electronics Engineers (IEEE) are being downloaded less than expected given their portfolio (Figure 3).

**Figure 3.**
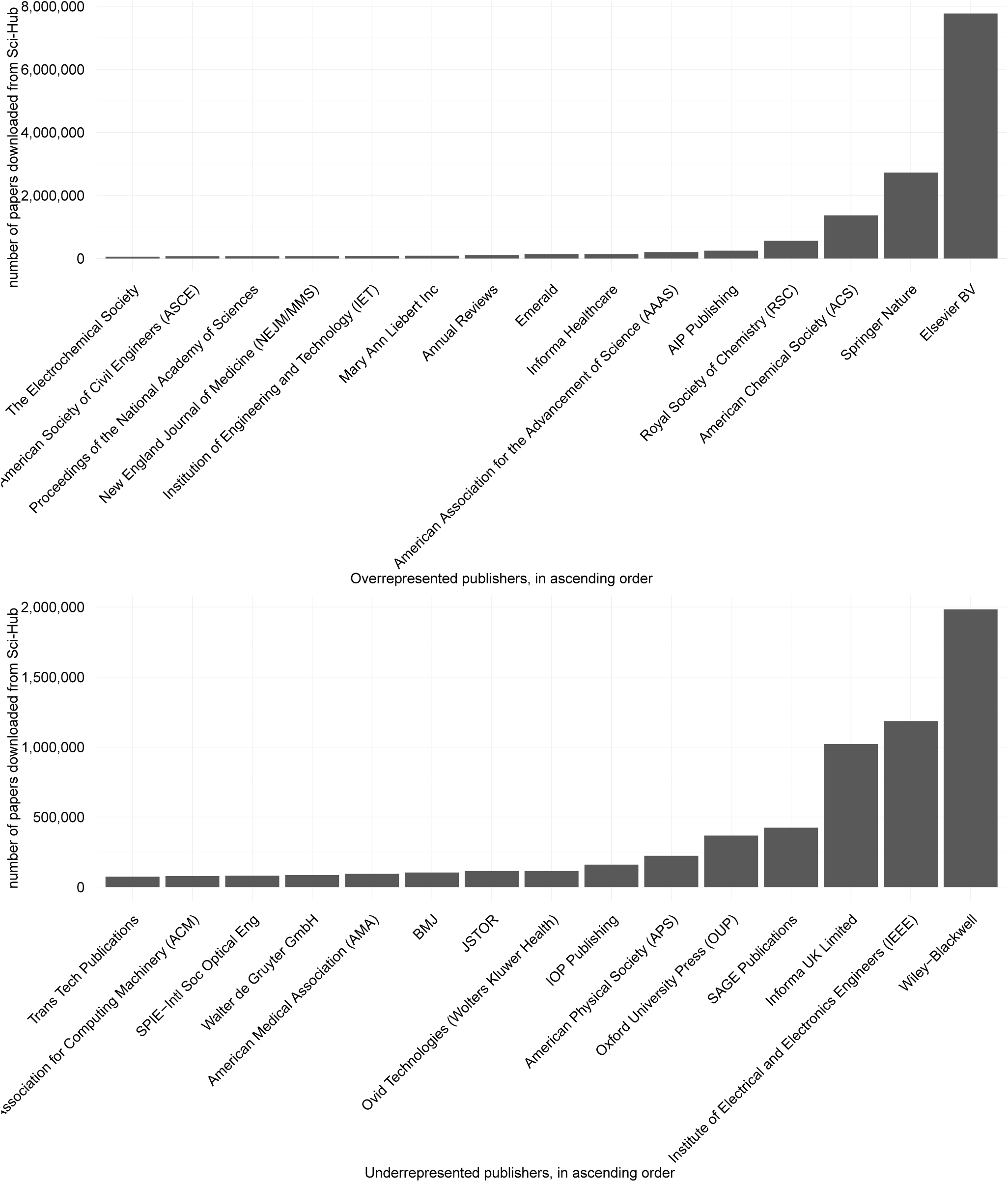
The most downloaded publishers that are either overrepresented (top) or underrepresented (bottom).

## Discussion

Earlier investigations into the data provided through *Sci-Hub* and *LibGen* focused large on either on the material being accessed [13] or on the data stored in these resources [10]. This study is the first to make use of both the whole corpus of *Sci-Hub* as well as data on how this corpus is being accessed by its users.

### Why *Sci-Hub?*

Comparing actual usage with the background set of articles shows that articles from recent history are highly sought for, giving some evidence that embargoes prior to making publications Open Access seem to become less effective. These findings are in line with prior research into the motivations for crowd-sourced, peer2peer academic file sharing [21]. While embargoes have impact on the use of those publications [22], these hurdles are being surpassed more and more surpassed *Black Open Access* [11], as provided by *Sci-Hub*.

While a good part of the literature available through *Sci-Hub* seems to be rarely accessed, the long tail of, publications, especially older ones, seems to be put to use ‐ albeit at a lower frequency. With DOIs that are unresolvable due to issues on publishers’ sides [23], and with Open Access publications that disappear behind accidental paywalls [24], this use for *Black Open Access* might play an important role and needs to be investigated more closely. It is worth noting that all analyses related to the number of downloads are limited to the six month period between September 2015 and February 2016, and do not necessarily reflect the complete use of *Sci-Hub*.

### Who’s reading?

Looking at the disproportionately frequented journals, one finds that 12 of the 20 most downloaded journals can broadly be classified as being within the subject area of chemistry. This is an effect that has also been seen in a prior study looking at the downloads done from *Sci-Hub* in the United States [12]. In addition, publishers with a focus on chemistry and engineering are also amongst the most highly accessed and overrepresented. While it is unclear whether this imbalance comes due to lack of access by university libraries, it’s noteworthy that both disciplines have a traditionally high number of graduates who go into industry. The 2013 *Survey of Doctorate Recipients* of the National Center for Science and Engineering Statistics (NCSES) of the United States finds that 50% of chemistry graduates and 58% of engineering graduates move to private, for-profit industry while only 32% and 27% respectively stay at educational institutions [25]. In comparison, in the life sciences these numbers are nearly switched, with 52% of graduates staying at educational institutions, which presumably offer more access to the scientific literature.

### *Non solus*. Or at least not completely

The prior analysis of the roughly 28 million downloads done through *Sci-Hub* showed a bleak picture when it came to the diversity of actors in the academic publishing space, with around 1/3 of all articles downloaded being published through Elsevier [13]. The analysis presented here puts this into perspective with the whole space of academic publishing available through *Sci-Hub*, in which Elsevier is also the dominant force with ∼24% of the whole corpus. The general picture of a few publishers dominating the market, with around 50% of all publications being published through only 3 companies, is even more pronounced at the usage level compared to the complete corpus, perpetuating the trend of *the rich getting richer*. Only 11% of all publishers, amongst them already dominating companies, are downloaded more often than expected, while publications of 45% of all publishers are significantly less downloaded.

## Conclusions

The analyses presented here suggest that *Sci-Hub* is used for a variety of reasons, by different populations. While most usage is biased towards getting access to recent publications, there is a subset of users interested in getting historical academic literature. Compared to the complete corpus, *Sci-Hub* seems to be a convenient resource, especially for engineers and chemists, as the overrepresentation shows. Lastly, when it comes to the representation of publishers, the *Sci-Hub* data shows that the academic publishing field is even more of an oligopoly in terms of actual usage when compared to the amount of literature published. Further analysis of how, by whom and where *Sci-Hub* is used will undoubtedly shed more light onto the practice of academic publishing around the globe.

## Data availability

All the data used in this study, as well as the code to analyze the data and create the figures, is archived on Zenodo as *Data and Scripts for Looking into Pandora’s Box: The Content of Sci-Hub and its Usage* (DOI, 10.5281/zenodo.472493).

In addition the analysis code can also be found on GitHub at *http://www.github.com/gedankenstuecke/scihub*.

## Acknowledgements

The author wants to acknowledge Alexandra Elbakyan, for releasing both data sets used in this study. Further thanks go to John Bohannon, who analyzed and helped in releasing the initial data on downloads from *Sci-Hub*. Further thanks go to Athina Tzovara and Philipp Bayer, for fruitful discussion of this manuscript as well as the statistics and analyses involved.

## supplementary-material

**Figure S1.**
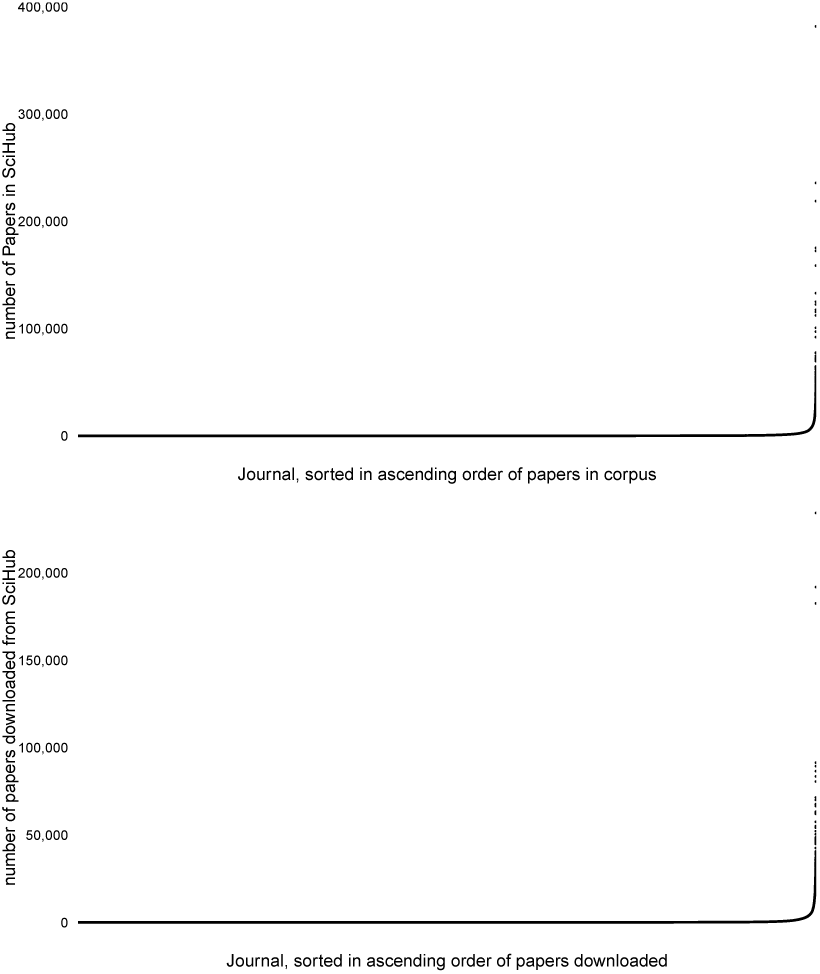
Top: The distribution of publications per journal in the whole corpus, sorted in ascending order of articles. Bottom: The distribution of downloads per journals, sorted in ascending order of downloads.

**Figure S2.**
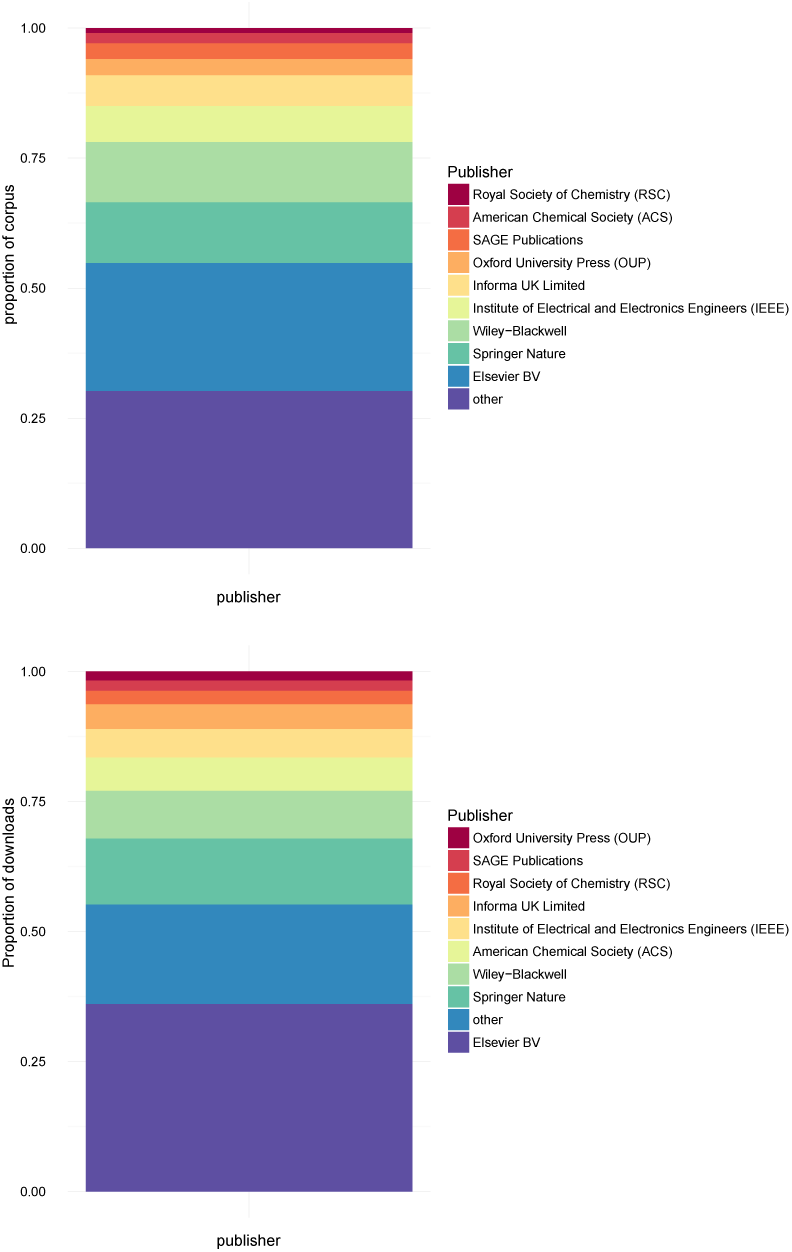
The proportion of the whole content as aggregated by publisher, both for the corpus (top) and downloads (bottom). Sorted by number of publications in the respective dataset. Only the 9 most frequent publishers are listed, smaller ones are grouped as *other*.

## References

[1] Bodó Balázs. Pirates in the library – an inquiry into the guerilla open access movement. Paper prepared for the 8th Annual Workshop of the International Society for the History and Theory of Intellectual Property, CREATe, University of Glasgow, UK, July 6-8, 2016, 2016. URL https://ssrn.com/abstract=2816925.

[2] Vincent Larivière, Stefanie Haustein, and Philippe Mongeon. The oligopoly of academic publishers in the digital era. PLOS ONE, 10(6):e0127502, 2015. doi: 10.1371/journal.pone.0127502.

[3] Paul Royster. A brief history of open access (accessed 4th of april, 2017), 2016. URL http://bit.ly/2nAgxKI.

[4] Jonathan P. Tennant, Francois Waldner, Damien C. Jacques, Paola Masuzzo, Lauren B. Collister, and Chris. H. J. Hartgerink. The academic, economic and societal impacts of open access: an evidence-based review. F1000Research, 5:632, 2016. doi: 10.12688/f1000research.8460.3.

[5] Diana Wildschut. The need for citizen science in the transition to a sustainable peer-to-peer-society. Futures, 2017. doi: 10.1016/j.futures.2016.11.010.

[6] Declan Butler. Gates foundation announces open-access publishing venture. Nature, 543(7647):599–599, 2017. doi: 10.1038/nature.2017.21700.

[7] Najko Jahn and Marco Tullney. A study of institutional spending on open access publication fees in germany. PeerJ, 4:e2323, 2016. doi: 10.7717/peerj.2323.

[8] Jim Giles. Trust gives warm welcome to open access. Nature, 432(7014):134–134, 2004. doi: 10.1038/432134a.

[9] Madian Khabsa and C. Lee Giles. The number of scholarly documents on the public web. PLoS ONE, 9(5):e93949, 2014. doi: 10.1371/journal.pone.0093949.

[10] Guillaume Cabanac. Bibliogifts in libgen? a study of a text-sharing platform driven by biblioleaks and crowdsourcing.Journal of the Association for Information Science and Technology, 67(4):874–884, 2015. doi: 10.1002/asi.23445.

[11] Bo-Christer Björk. Gold, green, and black open access. Learned Publishing, 2017. doi: 10.1002/leap.1096.

[12] Gabriel J. Gardner, Stephen R. McLaughlin, and Andrew D. Asher. Shadow libraries and you: Sci-hub usage and the future of ill. In ACRL 2017, Baltimore, Maryland, March 22–25, 2017, 2017.

[13] John Bohannon. Who’s downloading pirated papers? everyone. Science, 2016. doi: 10.1126/science.aaf5664.

[14] Elsevier inc. et al v. sci-hub et al case no. 1:15-cv-04282-rws (2015).

[15] Simon Oxenham. Meet the robin hood of science (accessed 4th of april, 2017), 2016. URL http://bit.ly/1nZx4YQ.

[16] J Bohannon and A Elbakyan. Data from: Who’s downloading pirated papers? everyone, 2016. URL http://dx.doi.org/10.5061/dryad.q447c.

[17] Bastian Greshake. Correlating the sci-hub data with world bank indicators and identifying academic use. The Winnower, 2016. doi: 10.15200/winn.146485.57797.

[18] Mark Hahnel. List of dois of papers collected by scihub. figshare, 2017. doi: 10.6084/m9.figshare.4765477.v1.

[19] Yoav Benjamini and Yosef Hochberg. Controlling the false discovery rate: A practical and powerful approach to multiple testing. Journal of the Royal Statistical Society. Series B (Methodological), 57(1):289–300, 1995.

[20] W. Snell. De cometarum materia, qui in solis vicinia non exarserunt. In Descriptio Cometæ, pages 53–57. Elsevier BV, 1619. doi: 10.1016/b978-1-4933-0406-6.50011-5.

[21] Carolyn Caffrey Gardner and Gabriel J. Gardner. Fast and furious (at publishers): The motivations behind crowd-sourced research sharing. College and Research Libraries, 78(2):131–149, 2017. doi: 10.5860/crl.78.2.131.

[22] Jim Ottaviani. The post-embargo open access citation advantage: It exists (probably), it’s modest (usually), and the rich get richer (of course). PLOS ONE, 11(10):e0165166, 2016. doi: 10.1371/journal.pone.0159614.

[23] Ross Mounce. Comparing oup to other publishers (accessed 4th of april, 2017), 2017. URL http://bit.ly/2nU6dko.

[24] Ross Mounce. Hybrid open access is unreliable (accessed 4th of april, 2017), February 2017. URL http://bit.ly/2ouDQdU.

[25] National Center for Science and Engineering Statistics. 2013 survey of doctorate recipients (accessed 4th of april, 2017), 2014. URL http://bit.ly/2oCuutw.

